# A 3D Structural Interactome to Explore the Impact of Evolutionary Divergence, Population Variation, and Small-molecule Drugs on SARS-CoV-2-Human Protein-Protein Interactions

**DOI:** 10.1101/2020.10.13.308676

**Authors:** Shayne D. Wierbowski, Siqi Liang, You Chen, Nicole M. Andre, Steven M. Lipkin, Gary R. Whittaker, Haiyuan Yu

**Author notes:** Correspondence should be addressed to H.Y.

## Abstract

The recent COVID-19 pandemic has sparked a global public health crisis. Vital to the development of informed treatments for this disease is a comprehensive understanding of the molecular interactions involved in disease pathology. One lens through which we can better understand this pathology is through the network of protein-protein interactions between its viral agent, SARS-CoV-2, and its human host. For instance, increased infectivity of SARS-CoV-2 compared to SARS-CoV can be explained by rapid evolution along the interface between the Spike protein and its human receptor (ACE2) leading to increased binding affinity. Sequence divergences that modulate other protein-protein interactions may further explain differences in transmission and virulence in this novel coronavirus. To facilitate these comparisons, we combined homology-based structural modeling with the ECLAIR pipeline for interface prediction at residue resolution, and molecular docking with PyRosetta. This enabled us to compile a novel 3D structural interactome meta-analysis for the published interactome network between SARS-CoV-2 and human. This resource includes docked structures for all interactions with protein structures, enrichment analysis of variation along interfaces, predicted ΔΔG between SARS-CoV and SARS-CoV-2 variants for each interaction, predicted impact of natural human population variation on binding affinity, and a further prioritized set of drug repurposing candidates predicted to overlap with protein interfaces^†^. All predictions are available online^†^ for easy access and are continually updated when new interactions are published.

† Some sections of this pre-print have been redacted to comply with current bioRxiv policy restricting the dissemination of purely in silico results predicting potential therapies for SARS-CoV-2 that have not undergone thorough peer-review. The results section titled “Prioritization of Candidate Inhibitors of SARS-CoV-2-Human Interactions Through Binding Site Comparison,” Figure 4, Supplemental Table 9, and all links to our web resource have been removed. Blank headers left in place to preserve structure and item numbering. Our full manuscript will be published in an appropriate journal following peer-review.

## Introduction

The ongoing global COVID-19 pandemic caused by the infection of SARS-CoV-2 has to date infected more than 30 million people and caused more than 940,000 deaths worldwide^1^. Coronaviruses are a family of enveloped viruses that cause respiratory and/or enteric tract infections in a wide range of avian and mammalian hosts^2^. To date, seven well characterized coronaviruses infect humans^3–5^ with severity ranging from a mild respiratory illness to severe pneumonia and acute respiratory distress syndrome (ARDS). Among these, SARS-CoV-2 is unique in its characteristics being both highly transmissible and capable of causing severe disease in a subset of individuals; whereas other human coronaviruses are either highly transmissible yet generally not highly pathogenic (e.g. HCoV-229E, HCoV-OC43) or highly pathogenic but poorly transmissible (SARS-CoV and MERS-CoV). SARS-CoV-2 is also unique in its range of infection and pathogenesis^6, 7^. While the vast majority of infected individuals (~25-35%) experience only mild or minimal symptoms, ~1-2% of infected patients die primarily from severe respiratory failure and acute respiratory distress syndrome^8, 9^. The differences in morbidity, hospitalization, and mortality among different ethnic groups—particularly Blacks and Hispanics^10–15^—could not be fully explained by cardiometabolic, socioeconomic, or behavioral factors, suggesting that human genetic variation may significantly impact SARS-CoV-2 pathogenicity. Therefore, insights into the evolution of SARS-CoV-2, its markedly elevated transmission rate relative to SARS-CoV, and dynamic range of symptoms are currently of key areas of interest. These traits are likely driven by differences in the mechanism of pathology and interactions between the virus and its host cells, but their specific causes are yet to be characterized.

One avenue to better understand the mechanisms of either viral- or bacterial-infection and pathology is through studying the network of protein-protein interactions that occur between a pathogen and its host. Viral-human interactome maps have previously been compiled for SARS-CoV^16^, HIV^17^, Ebola virus^18^, and Dengue and Zika viruses^19^ among others. Recently, an affinity-purification mass-spectrometry approach has been applied to 29 SARS-CoV-2 proteins uncovering 332 viral-human interactions^20^. These inter-species interactions play central roles in disease progression including, acting as facilitators of pathogen entry into host cells^21–26^, inducing an inhibitory effect on host response proteins and pathways^27–29^, and hijacking cell signaling or metabolism to accelerate cellular—and consequentially viral—replication^30–32^. Understanding the structure and dynamics of these interactions can provide critical insights into their effect on the cell. For instance, determination of the structure of an interaction between poxvirus chemokine inhibitor vCCI and human MIP-1β revealed that the viral-human binding interface occluded domains vital to chemokine homodimerization, receptor binding, and interactions with GAG, thus explaining how this interaction elicits an inhibitory effect on chemokine signaling^29^. Additionally, the dynamics of an interaction between a herpesvirus cyclin and human cdk2 revealed that although the binding interface was distinct from that between human cyclin A and cdk2, the net conformational impact on cdk2 effectively mimicked that of the native interaction leading to dysregulated cell cycle progression^31^.

Because protein-protein interactions are responsible for mediating the majority of protein function^33–35^, targeted disruption of these interactions by small molecule inhibitors that compete for the same binding site can offer a precise toolkit to modulate cellular function^33, 35–38^. For example, BCL-2 inhibitors that displace bound anti-apoptotic BCL-X interactors can treat chronic lymphocytic leukemia pathogenesis^39, 40^. Targeted disruption of protein interactions can be particularly effective in viral networks due to their small proteomes with highly optimized function, and potent inhibitors of key interactions have been developed against viral proteins. Targeted disruption of viral complexes— particularly those responsible for viral replication—has been successful in vaccinia virus^41^ and human papilloma virus therapies^42, 43^. In particular, disruption of viral-host protein-protein interactions explicitly involved in early viral infection is an important therapeutic strategy. Discovery that a population variant in the membrane protein CCR5 conferred resistance to HIV-1 by disrupting its interaction with the viral envelope glycoprotein led to the development of Maraviroc as an FDA approved treatment for HIV-1 that functions by blocking the interface for this interaction^23, 44^.

Here we apply a comprehensive interactome modeling framework to construct a 3D structural interactome between SARS-CoV-2 and human protein based on the current interactome map published by Gordon *et al*.^20^ Our framework consists of homology modeling to maximize structural coverage of the SARS-CoV-2 proteome, applying our previous ECLAIR classifier^45^ to identify interface residues for the whole SARS-CoV-2-human interactome, followed by atomic resolution interface modeling through guided docking in PyRosetta^46^. We additionally carried out in-silico scanning mutagenesis to predict the impact of mutations on interaction binding affinity and performed a comparison of protein-protein and protein-drug binding sites. We compile all results from our structural interactome into a user-friendly web server allowing for quick exploration of individual interactions or bulk download and analysis of the whole dataset. Further, we explore the utility of our interactome modeling approach in identifying key interactions undergoing evolution along viral protein interfaces, highlighting population variants on human interfaces that could modulate the strength of viral-host interactions to confer protection from or susceptibility to COVID-19, and prioritizing drug candidates predicted to bind competitively at viral-human interaction interfaces.

## Results

### Enrichment of divergence between SARS-CoV and SARS-CoV-2 at spike-ACE2 binding interface

To highlight the utility of computational and structural approaches to model the SARS-CoV-2-human interactome, we first examined the interaction between the SARS-CoV-2 spike protein (S) and human angiotensin-converting enzyme 2 (ACE2) (**Fig 1.a**). This interaction is key for viral entry into human cells^3^ and is the only viral-human interaction with solved crystal structures available in both SARS-CoV^47^ and SARS-CoV-2^48–50^. Comparison between SARS-CoV and SARS-CoV-2 revealed that sequence divergence of the S protein was highly enriched at the S-ACE2 interaction interface (**Fig 1.a**; Log2OddsRatio=2.82, p=1.97e-5), indicating functional evolution around this interaction. To explore the functional impact of these mutations on this interaction, we leveraged the Rosetta energy function^51^ to estimate the change in binding affinity (ΔΔG) between the SARS-CoV and SARS-CoV-2 versions of the S-ACE2 interaction (**Fig 1.b** and **1.c**). The predicted negative ΔΔG value of −14.66 Rosetta Energy Units (REU) indicates an increased binding affinity using the SARS-CoV-2 S protein driven by better optimized solvation and hydrogen bonding potential fulfillment along the ACE2 interface. Our result is consistent with the hypothesis that increased stability of the S-ACE2 interaction is one of the key reasons for elevated transmission of SARS-CoV-2^52^. Moreover, recent experimental energy kinetics assays have shown that SARS-CoV-2 S protein binds ACE2 with 10-20-fold higher affinity than that of SARS-CoV S protein^53^ supporting the conclusions from our computational modeling.

**Figure 1.**
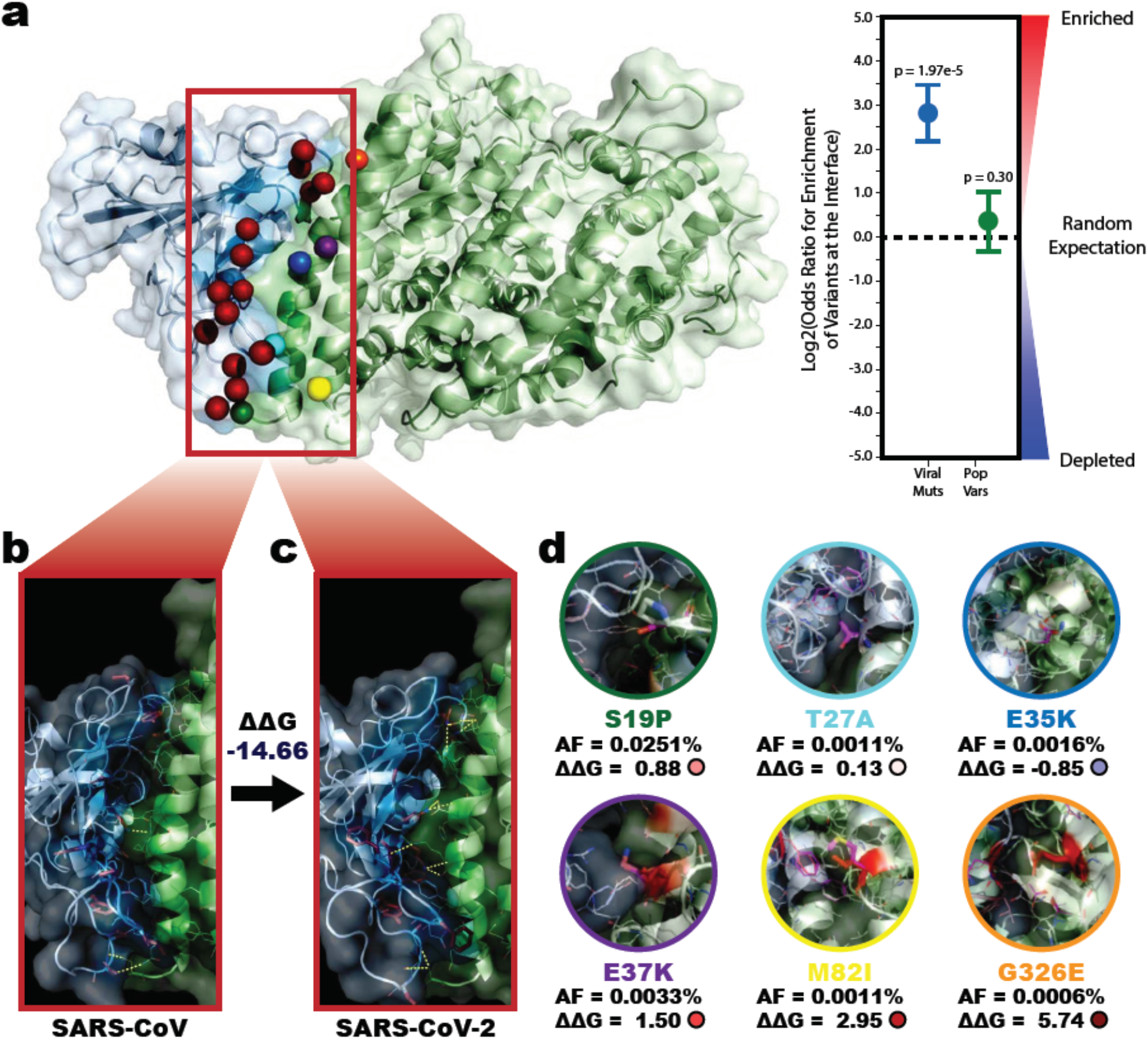
Enrichment and predicted impact of divergences between SARS-CoV and SARS-CoV-2 along the S-ACE2 interface. **a,** Co-crystal structure of the interaction between SARS-CoV-2 Spike protein (S) with human ACE2 (PDB 6LZG). All 15 sequence divergences between SARS-CoV and SARS-CoV-2 Spike protein interfaces are highlighted as red spheres while all 6 population variants on the ACE2 protein interface are highlighted as green (ACE2_S19P), cyan (ACE2_T27A), blue (ACE2_E35K), purple (ACE2_E37K), yellow (ACE2_M82I), and orange (ACE2_G326E) spheres. Enrichment of these variants on the interface are reported for SARS-CoV-2 (Log2OR=2.82, p=1.97e-5) and human (Log2OR=0.38, p=0.30) shown to the right. Error bars indicate ± SE. **b, c,** Zoomed in interface views for the SARS-CoV S-ACE2 structure (PDB 6CS2) and SARS-CoV-2 S-ACE2 structure (PDB 6LZG). Sequence divergences between the two Spike proteins are highlighted as red sticks. Inter-protein polar contacts that contribute to stabilizing the interaction are shown as yellow dashed lines. The binding energy (ΔG) of each interaction was estimated using PyRosetta and the change in this binding energy (ΔΔG) is reported. The negative value (ΔΔG=−14.66 Rosetta Energy Units (REU)) indicates the interaction is more stable (lower energy) in the SARS-CoV-2 version of the interaction. **d,** Comparison of the impact of each of the ACE2 population variants. Mutated structures containing the population variant are shown over the original structure (magenta). The mutated residue is shown as sticks. Residues whose predicted impact on binding energy was affected are colored from blue (decreased ΔΔG) to white (no change) to red (increased ΔΔG). The gnomAD reported allele frequency and predicted ΔΔG for each mutation are reported. Outlines of each structures are consistent with the spheres in **a**.

Medical experts have also noted a wide range in severity of and susceptibility to SARS-CoV-2 among individuals^6, 7, 54^. Several hypotheses for genetic predisposition models have been proposed including that expression quantitative trait loci (eQTLs) may up- or down-regulate host response genes and that functional coding variants may alter viral-human interactions^55, 56^. For instance, a recent RNA-sequencing analysis suggested that higher expression of ACE2 in Asian males effectively provides more routes of entry for the virus and could explain increased susceptibility among this population^57^. Alternatively, missense population variants in ACE2 could modulate susceptibility to infection by strengthening or weakening the S-ACE2 interaction. A total of six missense population variants reported in gnomAD^58^ localize to the S-ACE2 interface. Using a mutation scanning pipeline in PyRosetta^59, 60^ we predicted the impact on binding affinity for each variant (**Fig 1.d**). The three population variants predicted to have the largest impact on S-ACE2 binding affinity— ACE2_E37K (ΔΔG=1.50), ACE2_M82I (ΔΔG=2.95), and ACE2_G326E (ΔΔG=5.74)—were all consistent with previous experimental screens which identified them as putative protective variants exhibiting decreased binding of ACE2 to S^61, 62^. Cumulatively, our results highlight the usefulness of a 3D structural interactome modeling approach in identifying interactions and mutations important for viral infection, pathogenesis, and transmission.

### Constructing the 3D Structural SARS-CoV-2-Human Interactome

After successfully applying our modeling approaches to recapitulate the effect of mutations along the S-ACE2 interface, we expanded our efforts to compile a complete 3D structural interactome between SARS-CoV-2 and human proteins. An early interactome screen by Gordon *et al*.^*20*^ uncovered 332 viral-human interactions that provide the foundation for our 3D interactome. To model these interactions, we first constructed homology models for 18 of 29 SARS-CoV-2 proteins with suitable templates (**Supplemental Figure 1**). Then, we predicted the interface residues for each interaction using our **<u>E</u>**nsemble **<u>C</u>**lassifier **<u>L</u>**earning **<u>A</u>**lgorithm to predict **<u>I</u>**nterface **<u>R</u>**esidues (ECLAIR) framework^45^. In total, our pipeline identified 692 interface residues across 18 SARS-CoV-2 proteins with an average 24.7 residues per interface and 6,763 across 190 human proteins with an average 20.3 residues per interface. A summary of the classifier utilization, prediction confidences, and interface size is provided in **Supplemental Figure 2**.

In order to add a structural component to our interactome map, and thereby enable modeling of the binding affinity for these interactions, we additionally performed docking in PyRosetta using our ECLAIR interface likelihood predictions to refine the search space (**Supplemental Figure 3**). Structural models were available for both the human and SARS-CoV-2 proteins in 250 out of 332 interactions (75%) making them amenable to guided docking experiments. After performing up to 50 independent docking experiments for each interaction and retaining the top-scored conformation we report 959 interface residues across 18 SARS-CoV-2 proteins with an average 18.2 residues per interface and 4,483 across 250 human proteins with an average 17.9 residues per interface (**Supplemental Figure 2.g**). For all analyses, interface annotations provided from docking experiments were prioritized over our ECLAIR predictions. The full interface annotations from our ECLAIR and docking predictions are available in **Supplemental Table 1** and **Supplemental Table 2** respectively.

### Perturbation of Human Proteins by Disease Mutations and Binding of SARS-CoV-2 Interactions Occur at Distinct Sites

After constructing the 3D interactome between SARS-CoV-2 and human, we first looked for evidence of interface-specific variation by mapping both gnomAD^58^ reported human population variants (**Supplemental Table 3**) and sequence divergences between SARS-CoV and SARS-CoV-2 (**Supplemental Table 4**) onto the predicted interfaces. In general, conserved residues have been shown to cluster at protein-protein interfaces^63^, and a recent analysis of SARS-CoV-2 structure and evolution likewise concluded that highly conserved surface residues were likely to drive protein-protein interactions^64^. Consistent with these prior findings at an interactome-wide level, we observed significant depletion for both viral and human variation along the predicted interfaces comparable to that observed on solved human-human interfaces (**Fig 2.a**).

**Figure 2.**
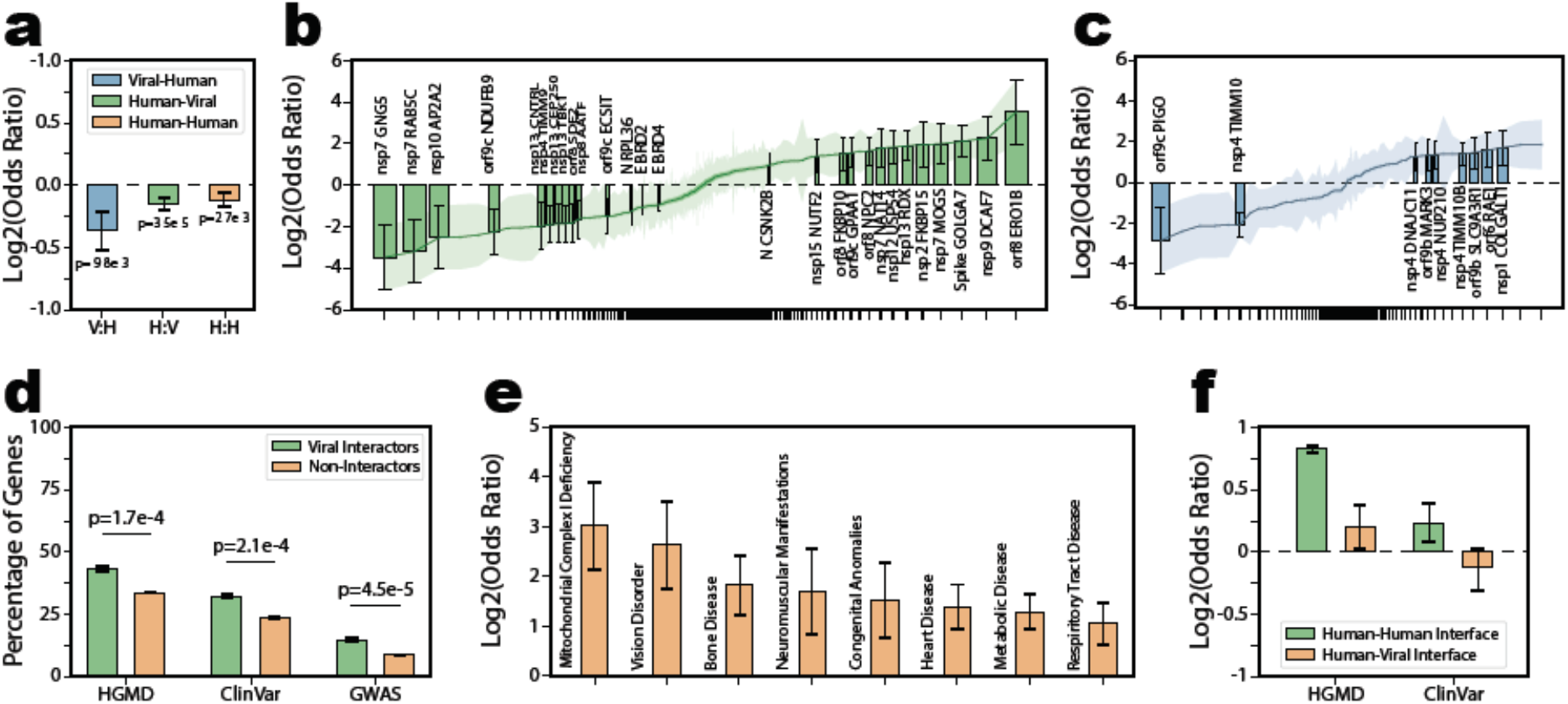
Enrichment of sequence divergences and disease mutations across all SARS-CoV-2-Human interaction interfaces. **a,** Overall enrichment for viral sequence divergence along viral-human (V:H, Log2OR=−0.37, p=9.76e-3) interfaces and human population variants along human-viral (H:V, Log2OR=−0.16, p=3.47e-5) or human-human (H:H, Log2OR=−0.13, p=2.67e-3) interfaces at an interactome level. Error bars indicate ± SE. **b, c,** Individual enrichment for human population variants and viral sequence divergences respectively on all SARS-CoV-2-human interaction interfaces. Interactions are sorted from most depleted to most enriched. The x-axis is skewed to compress non-significant interactions. Only interfaces with statistically significant enrichment or depletion are shown as bars and labeled. The remainder are shown as Log2OR (line) and error bars (colored area). Error bars indicate ± SE. **d,** Comparison of the percentage of human genes that interact with (green) or do not interact with (orange) SARS-CoV-2 that contain disease annotations in HGDM (Log2OR=0.57, p=1.70e-4), ClinVar (Log2OR=0.64, p=1.05e-4), and GWAS (Log2OR=0.89, p=4.54e-5) respectively. Genes targeted by SARS-CoV-2 proteins were significantly more likely to harbor disease mutations than non-interactors. Error bars indicate ± SE. **e,** A sample of individual disease terms enriched in human genes targeted by SARS-CoV-2. Full results are reported in **Supplemental Table 6**. Error bars indicate ± SE. **f,** Comparison of the enrichment of HGDM, ClinVar, and GWAS annotated mutations on human-vial interfaces or human-human interfaces for the same gene set. Although disease mutations were enriched on human-human interfaces (HGMD, Log2OR=0.82, p<1e-20; ClinVar, Log2OR=−0.13, p=0.24), no enrichment was observed on human-viral interfaces (HGMD, Log2OR=0.21, p=0.13; ClinVar, Log2OR=0.51, p<e-20). The GWAS category was removed from this analysis because most lead GWAS SNPs occurred in non-coding regions. Error bars indicate ± SE.

Nonetheless, considering each interaction individually, our analysis uncovered a 13 interaction interfaces enriched for human population variants (**Fig 2.b**), and 7 enriched for recent viral sequence divergences (**Fig 2.c**). A breakdown of variant enrichment on each interface is provide in **Supplemental Table 5**. The individual viral interfaces showing an unexpected degree of variation may—like the previously discussed S-ACE2 interface—be indicative of recent functional evolution around the viral-human interaction. Considering the slower rate of evolution in humans, enrichment of population variants along the human interfaces is unlikely to be a selective response to the virus. Rather, these interfaces with high population variation along the interfaces may represent edges in the interactome whose strength may fluctuate among individuals or between populations. Alternatively, enrichment and depletion of variation along the human-viral interfaces could help distinguish viral proteins that bind along existing—and therefore conserved—human-human interfaces from those that bind using novel interfaces—that would be less likely to be under selective pressure.

To further explore the functional impact of naturally occurring variants on the human interactors of SARS-CoV-2, we considered variants with phenotypic associations as reported in HGMD^65^, ClinVar^66^ or the NHGRI-EBI GWAS Catalog^67^. Interactors of SARS-CoV-2 were significantly more likely than the rest of the human proteome to harbor phenotypic variants in each of these databases (**Fig 2.d**). Notably, among the individual disease categories enriched in this gene set, several were consistent with reported comorbidities including heart disease, respiratory tract disease, and metabolic disease^68, 69^ (**Fig 2.e**; **Supplemental Table 6**). Disruption of native protein-protein interactions is one mechanism of disease pathology, and disease mutations are known to be enriched along protein interfaces^70, 71^. Human population variants on the predicted human-viral interface were more likely to be annotated as deleterious by SIFT^72^ and PolyPhen^73^ but showed identical allele frequency distributions compared to those off the interfaces (**Supplemental Figure 4**). However, mapping annotated disease mutations onto the protein interfaces only revealed significant enrichment along known human-human interfaces; no such enrichment was found on human-viral interfaces (**Fig 2.f**). This is likely because unlike with human-human interactions, mutations disrupting human-viral interactions would not disrupt natural cell function, and therefore would be unlikely to be pathogenic. Our finding that disease mutations and viral proteins affect human proteins at distinct sites is consistent with a two-hit hypothesis of comorbidities whereby proteins whose function is already affected by genetic background may be further compromised by viral infection.

### Analysis of Binding Affinity Changes Between SARS-CoV and SARS-CoV-2 Helps Identify Key Interactions

We next sought to explore the impact of sequence divergence in SARS-CoV-2 relative to SARS-CoV on viral-human interactions. Mutations between the two viruses were identified by pairwise alignment and the impacts of these mutations on the binding energy (ΔΔG) for 250 interactions amenable to docking were predicted using a PyRosetta pipeline^46, 59, 60^. Although the binding energy for most interactions was unchanged—either because no mutations occurred near the interface or because the mutations that did had marginal effect—we observed an increased likelihood of the divergence from SARS-CoV to SARS-CoV-2 resulting in decreased binding energy (i.e. more stable interaction) (**Fig 3.a**; **Supplemental Table 7**). The significant outliers in these ΔΔG predictions can help pinpoint key differences between the viral-human interactomes of SARS-CoV and SARS-CoV-2. We further note a wide range of affinity impacts among various human interactors of a single viral protein (**Fig 3.d**) and hypothesize that these differences may help identify the most important interactions.

**Figure 3.**
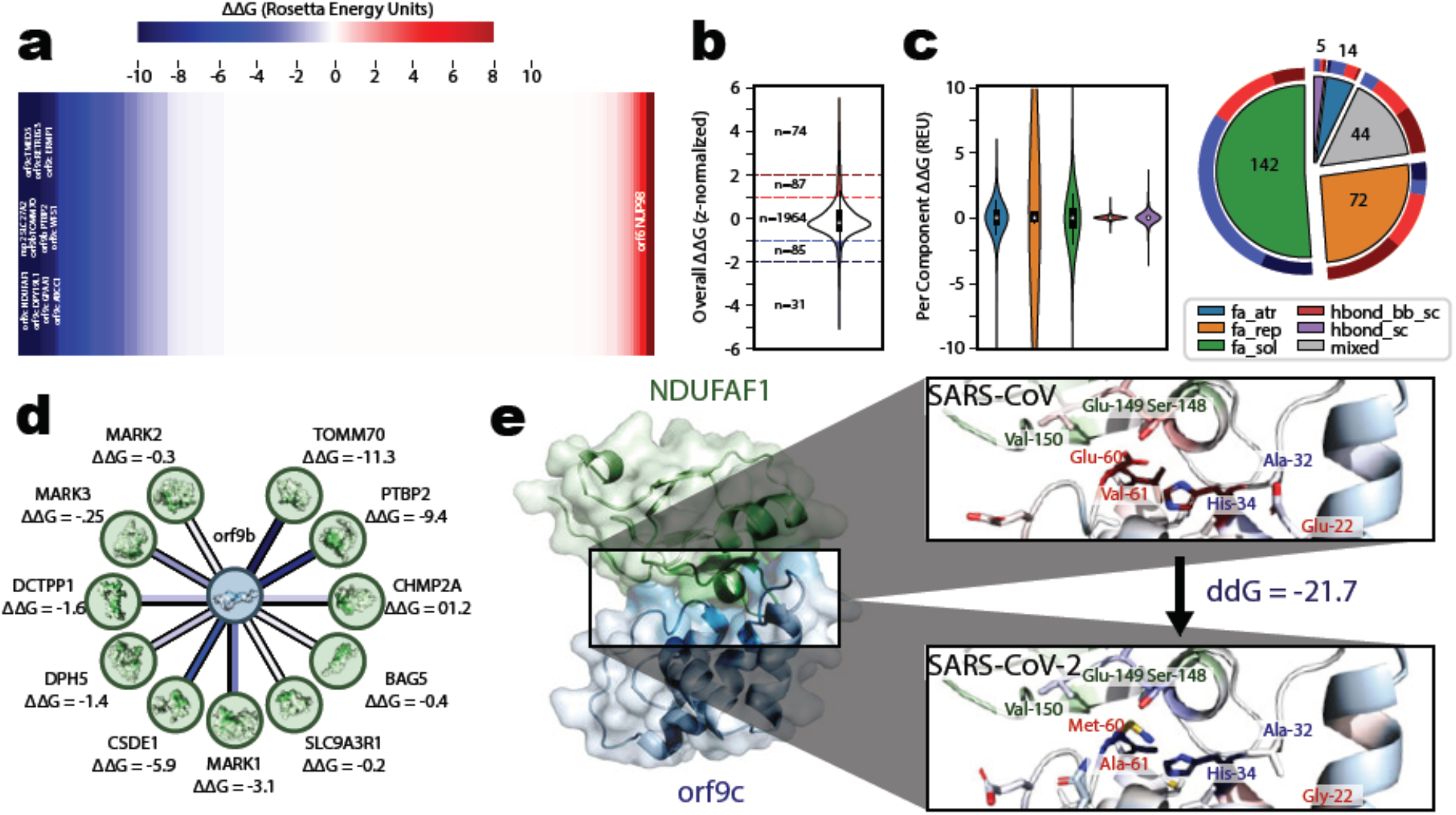
Predicted impact of sequence divergences on the binding affinity of SARS-CoV-2-Human interactions. **a,** Predicted changes in binding affinity from sequence divergences between SARS-CoV and SARS-CoV-2 on SARS-CoV-2-human interactions were predicted in PyRosetta. An overall representation of these ΔΔG predictions is reported (mean=−1.40 REU, std=6.16 REU) with interactions sorted from those with the largest decrease in binding energy (most stabilized relative to SARS-CoV) to those with the largest increase in binding energy (most destabilized relative to SARS-CoV). Outlier interactions with the change in binding energy at least one standard deviation from the mean are labeled. **b,** Distribution of the predicted binding affinity change from all human population variants on a SARS-CoV-2-human interface. Values were z-score normalized across each interface. Interface binding energy hotspots were binned as strongly disruptive (z-score ≥ 2, n=74), disruptive (1 ≤ z-score < 2, n=87), stabilizing (− 2 < z-score ≤ −1, n=85), or strongly stabilizing (z-score ≤ −2, n=31). All other population variants (−1 < z-score < 1, n=1,964) showed minimal impact of binding affinity. **c,** Breakdown of the contribution of each term in the PyRosetta energy function used for in-silico scanning mutagenesis for all population variants. Distributions for each term are shown on the left. A breakdown of which term contributed most heavily to the classification of all 277 interface hotspot population variants is shown on the right. **d,** Individual SARS-CoV-2-human interactions involving the same viral protein can have distinct interfaces with distinct predicted changes in binding affinity between SARS-CoV and SARS-CoV-2 versions of the protein. An example involving orf9b is highlighted where some interactions (e.g. TOMM70 and PTBP2) are predicted to be more stabilized in SARS-CoV-2 whereas others (e.g. BAG5, SLC9A3R1, and MARK2) are predicted to me unaffected. **e,** Docked structure for the interaction between SARS-CoV-2 orf9c and human NDUFAF1, alongside comparisons of the predicted interface using SARS-CoV (top) or SARS-CoV-2 (bottom) orf9c. Interface residues are colored by their predicted energy contribution from blue (stabilizing) to white (no impact) to red (destabilizing). Residues that differ between SARS-CoV and SARS-CoV-2 are labeled in red, while other residues with a major contribution to the binding affinity are labeled in green (NDUFAF1) or blue (orf9c). The overall predicted change in binding energy (ΔΔG=−21.7 REU) suggests the interaction is more stable (lower energy) in the SARS-CoV-2 version of the interaction.

To further explore the significance of these changes in interaction affinity, we considered those interactions with the largest decrease in binding energy; corresponding to the largest predicted increase in affinity. Specifically, we highlight the interaction between coronavirus orf9c and human mitochondrial NADH Dehydrogenase (Ubiquinone) 1 Alpha Subcomplex, Assembly Factor 1 (NDUFAF1) (**Fig 3.e**; ΔΔG=−21.7 REU). Two mutations contribute to the increased stability of this interaction in SARS-CoV-2. During the transition from the unbound to bound states of orf9c, His-34 rotates inwards to accommodate NDUFAF1 resulting steric tensions with orf9c residue 61. The mutation from the bulkier Val-61 (SARS-CoV) to Ala-61 (SARS-CoV-2) helps alleviate this tension resulting in overall more favorable energy state transition. Second, the mutation from the polar Glu-60 (SARS-CoV) to the nonpolar Met-60 (SARS-CoV-2) contributes to overall better solvation energy along the interface involving residue 60 of orf9c and the Ser-148, Glu-149, Val-150 stretch of NDUFAF1.

Although the precise role of coronavirus orf9c (sometimes annotated as orf14) has not yet been characterized, experiments in SARS-CoV have shown that it localizes to vesicular components^74^ and interactome data from SARS-CoV-2 reveal that it targets host mitochondrial proteins and could impact proteins responsible for modulating IkB kinase and NF-kB signaling pathways^20^. NDUFAF1 is a chaperone protein involved in the assembly of the mitochondrial complex I^75, 76^. RNAi screens have associated knockdown of NDUFAF1 with increased vaccinia virus infection^77^. Moreover, properly assembled mitochondrial complex I—including NDUFAF1—has been shown to be indispensable for reactive oxygen species based signaling to trigger expression of interleukins 2 and 4 and activate signaling of TLR7^78^, a SARS-CoV-2 ARDS predisposition gene^79^. Recent reports have suggested that the virus may hijack host metabolism through complex I to increase replication^80^. Additionally, NDUFAF1 harbors disease mutations for complex I deficiency and hypertrophic cardiomyopathy that may be linked to SARS-CoV-2 comorbidities. Our result predicting increased stability of the interaction between orf9c and NDUFAF1 in SARS-CoV-2 relative to SARS-CoV suggests a stronger impact on complex I assembly and potential disruption of downstream pathways.

We further identified several interactions with significantly altered binding energy in SARS-CoV-2 with potential links to cellular response to viral infection. In orf9c we additionally predicted increased affinity with a Golgi signaling protein, TMED5 (ΔΔG=−21 REU). Phenotypic screens have linked TMED5 to altered viral infectivity^77, 81, 82^ and decreased interleukin 8 secretion^83^. Although no known mechanisms directly link TMED5 to immune response, one of its interactors, TMED2^84^, is known to potentiate interferon response to viral infection^85^. Interaction with TMED5 may additionally be linked to viral transport within and secretion from its host^86^. Finally, the mitochondrial protein TOMM70 was predicted to bind orf9b with greater affinity (ΔΔG=−11 REU) in SARS-CoV-2. TOMM70 has been linked to mitochondrial antiviral signaling through downstream activation of interferon regulatory factors^87^.

### Predicting the Impact of Human Populations Variants on SARS-CoV-2-Human Protein Interactions

A dynamic range of patient responses and symptoms have been reported for SARS-CoV-2 infection. In previously studied viruses including HIV, underlying genetic variation can explain up to 15% of variation in patient response and overall viral load^88^. Moreover, previous high-throughput experiments suggest that, up to 10.5% of missense population variants can disrupt native protein-protein interactions^89^. Therefore, we hypothesize that some fraction of patient response to SARS-CoV-2 can be explained by missense variations and their impact on viral-human interactions. To explore this hypothesis, we performed in-silico scanning mutagenesis along all docked interfaces in PyRosetta. We identified as binding energy hotspot mutations all mutations with a predicted ΔΔG at least one standard deviation away from the mean for identical amino acid substitutions across the rest of the interface. In total, out of 2,241 population variants on eligible interfaces, 161 (7.2%) were identified as hotspots predicted to disrupt interaction stability, and 116 (5.2%) were identified as hotspots predicted to contribute to interaction stability (**Fig 3.b**). Most of the hotspot mutations were predicted to be driven by solvation or repulsive forces, with disruptive hotspots primarily being driven by repulsive forces and stabilizing hotspots primarily being driven by solvation forces (**Fig 3.c**). Results summarizing the predicted impact of all 2,241 population variants on the docked interfaces are provided in **Supplemental Table 8**.

### Prioritization of Candidate Inhibitors of SARS-CoV-2-Human Interactions Through Binding Site Comparison

#### A Web Server to Present the SARS-CoV-2-Human 3D Structural Interactome

In order to present the results from our experiments as a comprehensive easy-access resource to the general public, we constructed the SARS-CoV-2 human structural interactome web. All structures used for modeling, interface predictions, raw docking outputs, mutation binding impacts, and analyses described herein are directly available for download through our downloads page. Our homepage allows users quick navigation through the reported interactome to view results summarizing specific interactions of interest. Aside from providing a 3D view of the interaction and predicted interface residues, our results page provides four main functionalities (**Fig 5**).

**Figure 5.**
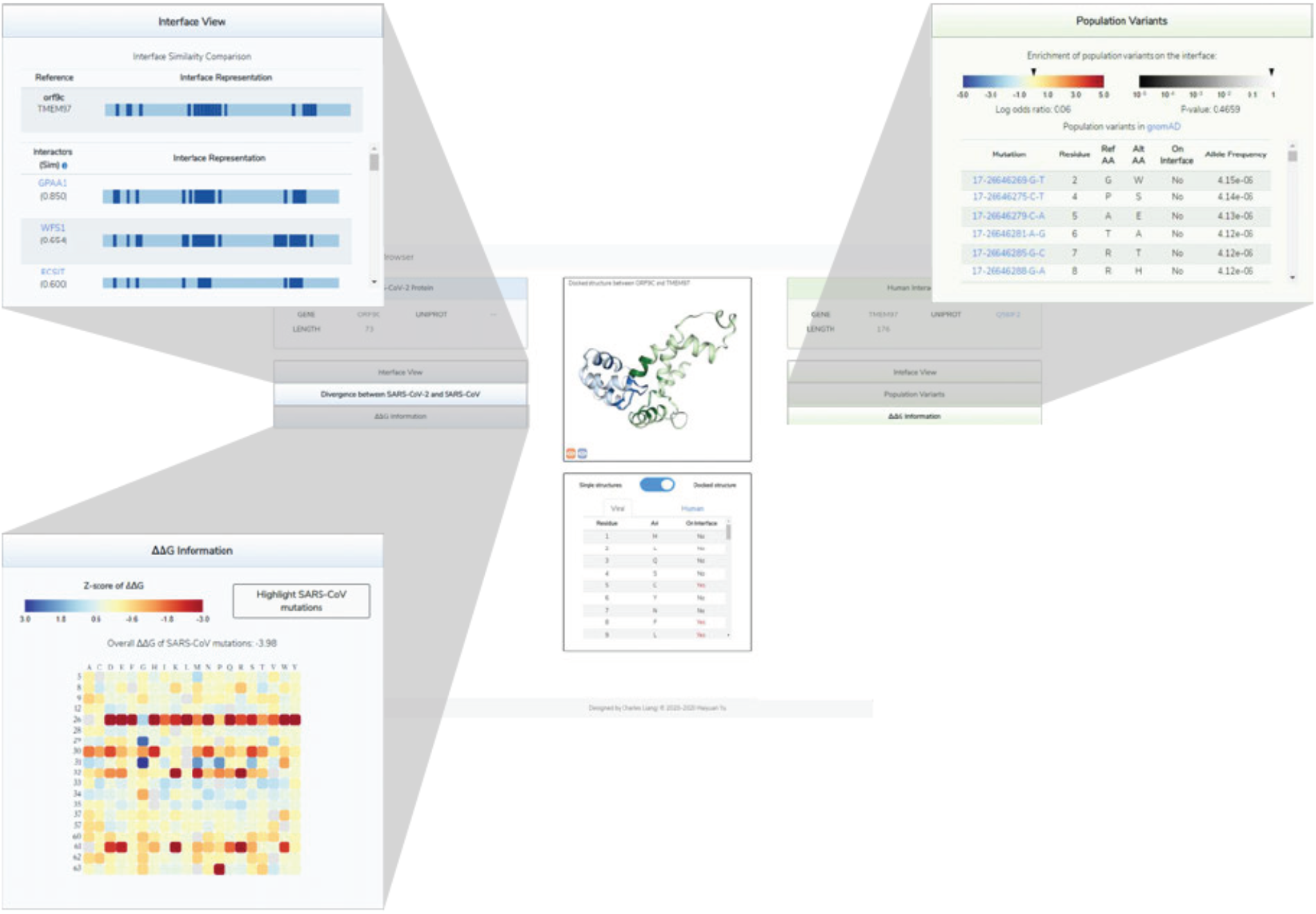
3D-SARS2 Structural Interactome Browser Overview. Overview of the main results page for exploring a given interaction in our 3D-SARS2 structural interactome browser. The main display contains information for both the SARS-CoV-2 and human proteins including structural displays for either the docked or single crystal structures as well as a table summarizing the interface residues for both proteins. Interface residues are colored dark blue and dark green for the viral and human proteins respectively. By default the page will display the docked structure if available. The display can be toggled between docked structures and single structures using the button in the bottom middle. When single structures display is selected residues will instead be colored based on the initial ECLAIR interface definition. Four categories of expandable panels containing additional analyses are provided. **upper left,** The interface view shows a linear representation of the protein sequence with interface residues annotated in dark blue or dark green. Interfaces for other interactors of the protein are shown underneath for easy comparison. **upper right,** The mutations panel summarizes either human population variants or viral sequence divergences on the protein. Mutations on the interface are labeled. **lower left,** The ΔΔG information panel summarizes the results from in-silico mutagenesis scanning along the interface. Results for each mutation are z-score normalized relative to the rest of the interface and colored on a blue (negative ΔΔG, stabilizing) to yellow (minimal impact) to red (positive ΔΔG, destabilizing). The heatmap can be filtered to only show values corresponding to known mutations on the interface.

The interface comparison panel (**Fig 5 top left**) provides a linear representation of protein sequence with interface annotations. Linear representations of all other known or predicted interfaces from the same protein are shown for comparison along with navigation links to explore other interaction partners either within the site (for viral-human interactions) or through InteractomeINSIDER^45^ (for human-human interactions). In particular, from the human perspective, this comparison may reveal biologically meaningful insights about the interface overlap and possible competition between viral and human interactors.

The mutations panel (**Fig 5 top right**) provides information on variation in each interaction partner. For the viral side, mutations are reported based on divergence from the SARS-CoV version of the protein. For the human side, all known population variants as reported in gnomAD are listed^58^. Additionally, we provide a log odds ratio describing the enrichment or depletion of variation along the interface. These results may help highlight interactions undergoing functional evolution along the viral interface.

For each interaction amenable to docking, the ΔΔG Information panel (**Fig 5 lower left**) compiles the results from our in-silico scanning mutagenesis to report binding affinity predictions for all possible mutations across the docked interface. Individual mutations and colored by their z-score normalized ΔΔG prediction. The results can be toggled to only show the impacts of known variants. On the viral side, a cumulative ΔΔG value is provided describing the predicted change in binding affinity between the SARS-CoV and SARS-CoV-2 versions of the protein.

The current version of the SARS-CoV-2 human structural interactome web server describes 332 viral-human interactions reported by Gordon *et al*.^20^. We will continue support for the web server with periodic updates as additional interactome screens between SARS-CoV-2 and human are published. As we update, a navigation option to select between the current or previous stable releases of the web server will be provided.

## Discussion

Overall, we present a comprehensive resource to explore the SARS-CoV-2-human protein-protein interactome map in a structural context. Analysis through this framework allows us to consider the recent evolution of SARS-CoV-2 in the context of its interactome map and to prioritize for further functional characterization key interactions. Likewise, our consideration of underlying variation in the human proteins that interact with SARS-CoV-2 may be valuable in explaining differences in response to infection. We particularly note that our observation that perturbation from underlying disease mutations and viral protein binding occur at distinct sites on the protein is of clinical interest. Further investigation into the combined role of these two sources of perturbation to better understand the mechanisms linked to comorbidities is warranted.

However, our work is not without limitation. Firstly, we note that although structural coverage from our homology modelling of SARS-CoV-2 proteins was robust (**Supplemental Figure 1**), the same could not be universally said of the human proteins. Although guided molecular docking was always done to orient the most likely interface residues on each structure towards each other, protein-protein docking using incomplete protein models introduces some bias and low coverage may exclude some true interface residues. For this reason, the initial ECLAIR interface annotations—which are less subject to structural coverage limitations—may provide orthogonal value. We additionally note that direct quantitative interpretation of predicted ΔΔG values using the Rosetta scoring function is often difficult since different term weights can be used in different setups. Although the same scoring function was used for all predictions described here, the relative magnitude of each term may change based on the size and composition of different proteins between interactions. For these reasons, we only employ a relative qualitative comparison of similar predictions when interpreting our scanning mutagenesis results. Moreover, the structure optimization after each mutation is applied focuses on side-chain repacking, therefore, our results focus only on mutations at or near the interface where the impact of side-chain repacking could be measured. We expect there may be some mutations with significant impact on binding affinity that act through refolding or other allosteric effects that are missed by our method.

Perhaps most importantly, we emphasize the importance of further experimental characterization to confirm the predictions made here. Nonetheless we believe our 3D Structural SARS-CoV-2-human Interactome web server will prove to be a key resource in informing hypothesis driven exploration of the mechanisms of SARS-CoV-2 pathology and host response. The scope, and potential impacts of our webserver will continue to grow as we incorporate the results of ongoing and future interactome screens between SARS-CoV-2 and human. Additionally, we note that our 3D structural interactome framework can be rapidly deployed to analyze future viruses.

## Online Methods

### Generation of Homology Models for SARS-CoV-2 Proteome

Homology-based modeling of all 29 SARS-CoV-2 proteins was performed in Modeller^90^ using a multiple template modeling procedure. In brief, a list of candidate template structures for each protein to be modelled was obtained by running BLAST^91^ against a reference containing all sequences in the Protein Data Bank (PDB)^92^. Templates were filtered to only retain those with at least 30% identify to the protein to be modelled, and the remaining templates were ranked using a weighted combination of percent identity and coverage as described previously^93^. To compile the final set of overlapping templates for modeling, first the top ranked template was selected as a seed. Overlapping templates were iteratively added to the set so long as 1) the new template increased the overall coverage by at least 10%, and 2) the new template retained a total percent identity no more than 25% worse than the initial seed template. Pairwise alignments between the protein to be modelled and the template set were generated using a Modeller alignment object with default settings. Alignments were manually trimmed to remove any regions with large gaps (at least 5 gaps in the alignment in a 10 residue window). Finally, alignment was carried out using the Modeller automodel function.

This approach generated suitable homology models for 18 out of 29 proteins. A visual representation of each structure and templates used is provided in **Supplemental Figure 1**.

### Phase One Interface Prediction Using ECLAIR

Interface predictions for all 332 interactions reported by Gordon *et al.*^20^ were made in two phases. In phase one, we leveraged our previously validated **<u>E</u>**nsemble **<u>C</u>**lassifier **<u>L</u>**earning **<u>A</u>**lgorithm to predict **<u>I</u>**nterface **<u>R</u>**esidues (ECLAIR)^45^ to perform an initial prediction of likely interface residues across all interactions. ECLAIR compiles five sets of features; biophysical, conservation, coevolution, structural, and docking. In brief, biophysical features are compiled using a windowed average of several ExPASy ProtScales^94^, conservation features are derived from the Jensen-Shannon divergence^95, 96^ across all available homologs for each protein, coevolution features between interacting proteins are derived from direct coupling analysis (DCA)^97^ and statistical coupling analysis (SCA)^98^ among paired homologs, structural features are obtained by calculating the solvent accessible surface area of available PDB^92^ or ModBase^99^ models using NACCESS^100^, and docking features are derived from an average of docking predictions made using Zdock^101^.

To accommodate predictions between SARS-CoV-2 and human, slight alterations were made. First calculation of coevolution features was impossible because the calculation of SCA and DCA requires analysis of multiple sequence alignments from paired homologs of both interacting proteins across at least 50 species. Because we here consider inter-species interaction, no one species could contain homologs of both the viral and human proteins. Second, the calculation of conservation features for the viral proteins were modified to account for conservation between both related viral species and various strains that have been sequenced in a single species. We typically only include one homolog per species in these calculations, but expanded our criteria here because availability of the protein conservation feature is a requirement for all of our higher-confidence classifiers. Finally, the calculation of structural features for the viral protein were overruled to use the manually provided homology models instead of pulling structures from the PDB or ModBase. A visual summary of the ECLAIR interface predictions is presented in **Supplemental Figure 2** and the initial prediction results are provided in **Supplemental Table 1**.

### Phase Two Interface Prediction Using Guided PyRosetta Docking

Interface predictions for all 332 interactions reported by Gordon *et al.*^20^ were made in two phases. In phase two, we generated atomic resolution models of 250 interactions by leveraging the Rosetta scoring function^51^ and prior probabilities obtained from ECLAIR predictions to perform guided docking. The remaining 82 interactions were missing reliable 3D models for at least one of their members and therefore were not amenable to docking. A schematic summary of our docking methodology is presented in **Supplemental Figure 3**.

Using the predicted interface probabilities reported by ECLAIR, we set up the initial docking conformation to explore a restricted search space for each docking simulation. In cases where multiple structures were available for the human protein, all structures were weighted based on the ECLAIR scores for the covered residues in each structure to maximize both coverage age inclusion of likely interface residues. For each protein in the interaction, we performed a linear regression classification in scikit-learn^102^ to optimally separate the likely interface residues from likely non-interface residues. The plane defined by this linear regression served as a reference to orient the structures along the y-axis with their most probable interface sides facing each other. The two chains were centered at (0, 0) along the x-z plane and separated a distance of 5 Å along the y-axis. For each docking attempt, a series of random perturbations from these initial conformations were made to search the nearby space. First, the human protein was rotated up to 360° along the y-axis to allow full exploration of different rotations of the two interfaces relative to each other. Second, to apply some flexibility to the plane predicting the interface vs. non-interface sides of each protein, up to 30° of rotation along the x- and z-axis were allowed for both the viral and human proteins. Finally, a random translation up to 5 Å in magnitude was applied to the human protein along the x-z plane so that the docking could explore contact points other than the center of masses along these axes.

After initializing these guided starting conformations, docking was simulated in PyRosetta^46^ using a modified version of the protein-protein docking methodology provided by Gray 2006^103^. The initial demo (https://graylab.jhu.edu/pyrosetta/downloads/scripts/demo/D100Docking.py) takes two chains from a co-crystal structure, applies a random perturbation, and re-docks them. Because randomized initial orientation was already handled as described previously, these steps were removed from our docking runs. In brief, the protein models were converted to centroid representation, slid into contact using the “interchain_cen” scoring function, and converted back to full atom representation, before having their side-chains optimized using the predefined “docking” and “docking_min” scoring functions. Up to 50 iterations of this guided docking were performed for each interaction, and the docked conformation resulting in the lowest Rosetta energy score was retained. The final docked interface annotations are provided in **Supplemental Table 2**.

### Definition of Interface Residues

To annotate interface residues from atomic resolution docked models, we used a previously described and established definition for interface residues^45^. In brief, the solvent accessible surface area (SASA) for both bound and unbound docked structures was calculated using NACCESS.^100^ We define as interface residue, any residue that is both 1) at the surface of a protein (defined as ≥ 15% relative accessibility) and 2) in contact with the interacting chain (defined as any residue whose absolute accessibility decreased by ≥ 1.0 Å^2^).

### Compilation of Viral SARS-CoV to SARS-CoV-2 Mutations and Human Population Variants

For analysis of genetic variation that may impact the viral-human interactome, two sets of mutations were compiled; 1) viral mutations between SARS-CoV and SARS-CoV-2, and 2) human population variants on proteins shown to interact with viral proteins.

For viral mutations, variations between SARS-CoV and SARS-CoV-2 versions of each proteins were collected. The representative sequences used for the 16 proteins in the SARS-CoV proteome were taken from the Proteomes section of UniProt (ID UP000000354)^104, 105^. Sequences for 29 SARS-CoV-2 proteins were obtained from the annotations by Gordon *et al.*^20^ based on genbank accession MN985325^106, 107^. Notably, UniProt accessions for the SARS-CoV proteome report two sequences for the uncleaved ORF1a and ORF1a-b which correspond to NSP1 through NSP16 in SARS-CoV-2. Variations between SARS-CoV and SARS-CoV-2 were reported after pairwise Needleman Wench alignment^108, 109^ (using Blosum62 scoring matrix, gap open penalty of 10 and gap extension penalty of 0.5) between the corresponding protein sequences in each species. A total of 1,003 missense variants were detected among 23 SARS-CoV-2 proteins. No variations were reported for orf3b, orf8, or orf10 because no suitable alignment could be made with a SARS-CoV sequence. Additionally, orf7b, nsp3, and nsp16 were excluded from this set because they were not involved in any viral-human interactions. The full list of SARS-CoV-2 mutations is reported in **Supplemental Table 4**.

Human population variants in all 332 human proteins shown to interact with SARS-CoV-2 proteins were obtained from gnomAD^58^. Programmatic queries to fetch all variants for a given gene were performed using gnomAD’s graphQL API. For details on performing gnomAD queries in this manner see the gnomad-api github page (https://github.com/broadinstitute/gnomad-browser/tree/master/projects/gnomad-api). Population variants were filtered to only retain missense variants. In order to map these gnomAD DNA-level SNPs to equivalent protein-level UniProt annotations, we used the Ensembl Variant Effect Predictor (VEP)^110^. After all mapping was completed, all variants were parsed to make sure that the reported reference amino acid and position matched up with the UniProt sequence. Roughly 95.6% of cases matched and the remaining variants that could not reliably be mapped to UniProt coordinates were dropped from our dataset. In total 127,528 human population variants were curated. The full list of human population variants from GnomAD is reported in **Supplemental Table 3**.

### Log Odds Enrichment Calculations

In order to determine if viral mutations or human populations variants were enriched at predicted interaction interfaces, odds ratios were calculated as described previously^111^. All odds ratios were log2 transformed to maintain symmetry between enriched and depleted values. For this particular use case, the equation for the odds ratio was…

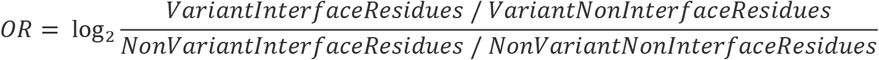

where “Variant” vs. “NonVariant” distinguishes positions in a protein sequence that respectively did and did not overlap with the set of variants, and “InterfaceResidues” vs. “NonInterfaceResidues” distinguishes positions in a protein sequence that respectively did and did not overlap with the predicted interface for a given interaction. To avoid arbitrary odds ratio inflation or depletion from missing data, in all cases where the interface residues were predicted by molecular docking, the odds ratio was altered to only account for positions that were included in the structural models used for docking. Enrichment calculations for disease mutations and overlap between drug and protein binding sites were calculated in the same manner with appropriately adjusted case and exposure categories.

### Curation of Disease Associated Variants

To explore whether human proteins interacting with SARS-CoV-2 proteins were enriched for disease or trait associated variants, three datasets were curated; the Human Gene Mutation Database (HGMD)^65^, ClinVar^66^, and the NHGRI-EBI GWAS Catalog^67^. Disease annotations from HGMD and ClinVar were obtained directly from their respective downloads pages and mapped to UniProt. For overall enrichment of individual disease terms among all human proteins interacting with SARS-CoV-2, disease terms were linked in an ontology based on the NCBI MedGen term relationships (https://ftp.ncbi.nlm.nih.gov/pub/medgen/MGREL.RRF.gz). When calculating enrichment, counts for each term were propagated through all parent nodes up to a singular root node. Significant terms were reported as the most general term with no more significant ancestor term (**Supplemental Table 6**, sheet 1). Raw enrichment values for all terms are also predicted (**Supplemental Table 6**, sheet 2).

For curation of disease and trait associations from NHGRI-EBI GWAS Catalog (http://www.ebi.ac.uk/gwas/)^67^, lead SNPs (p-value<5e-8) for all diseases/traits were retrieved on June 16, 2020. Proxy SNPs in high linkage disequilibrium (LD) (Parameters: R^2^ > 0.8; pop: “ALL”) for individual lead SNPs were obtained through programmatic queries to the LDproxy API^112^, which used phase 3 haplotype data from the 1000 Genomes Project as reference for calculating pairwise metrics of LD. Both lead SNPs and proxy SNPs were filtered to only retain missense variants.

Gene-level enrich calculations for disease / trait associated mutations in known interactors for SARS-CoV-2 and variant-level enrichment at the predicted interaction interfaces (either human-human interactions or human-viral interactions) were performed as described above.

### Estimation of ΔΔG from variation at the interfaces

In order to predict the impact of variation at viral-human interaction interfaces on binding affinity, two sets of ΔΔG predictions were made using PyRosetta^46^. First, to explore the overall binding energy contributions of each interface residue, and to predict the impact of all possible mutations along the interface, a scanning mutation ΔΔG approach was implemented based on protocols described previously^59, 60^. The implementation is essentially identical to a demo provided by the PyRosetta documentation (https://graylab.jhu.edu/pyrosetta/downloads/scripts/demo/D090Alascan.py) with one major exception: interface residues in the original demo are defined using an 8.0 Å inter-chain distance cutoff, which we overrule with our definition described above. In brief, we iterate over all interface residues for a given docked structure. For each position, the binding for the interaction from the wildtype structure is first estimated. The energy for the complex state is estimated following a PackRotamersMover optimization constrained to only adjust the side-chains of residues within 8.0 Å of the residue to be mutated. To estimate energy from the unbound state, the chains are first separated 500.0 Å to eliminate any interchain energy contributions, the structures are then optimized and scored identically to the complex state. The difference between these two values provides the binding energy for the wildtype structure.

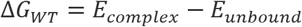

To estimate the binding energy for all 19 amino acid mutations possible at the given position, each mutation is made iteratively, and the ΔG_Mut_ is estimated identically to the wildtype except using the mutated structures. Finally, the overall impact on binding energy for each mutation is calculated as the difference between these two binding energies.

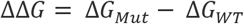

The scoring function used for these calculations is as described previously^59^ using the following weights; fa_atr=0.44, fa_rep=0.07, fa_sol=1.0, hbond_bb_sc=0.5, hbond_sc=1.0. To account for the stochasticity of the PackRotamersMover optimization between trials, all ΔΔG values are reported from an average of 10 independent trials which yielded overall low standard deviation between trials. To check whether an individual mutation’s average ΔΔG was significantly non-zero, a two-sided z-test between the 10 independent trials was performed. To account for average impact other same amino acid mutations at other positions along the interface, each average ΔΔG was z-normalized relative to the rest of the interface and outliers were called at ≥ 1 standard deviation away from the mean. Mutations that passed both criteria were identified as significant interface binding affinity hotspots.

Second, estimates of the overall impact of the cumulative set of mutations between SARS-CoV and SARS-CoV-2 were made using the same general framework. For each interaction, the binding energy for the SARS-CoV-2 version of the interaction was estimated identically to the wildtype binding energy. The binding energy for the SARS-CoV version of the interaction was estimated after applying all mutations between the two viruses (for simplicity, only amino acid substitutions were applied, a minority of mutations that comprised insertions or deletions could not be modelled). The only difference compared to the single mutation ΔΔG calculation was that multiple mutations were applied at once, and consequentially, the interface packing was done allowing side-chain rotamer optimization for all residues within 8.0 Å of any of the mutated residues. The ΔΔG values were calculated such that an interaction predicted to be more stable (lower binding energy) in the SARS-CoV-2 version of the interaction compared to the SARS-CoV version of the interaction would have a negative ΔΔG.

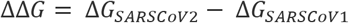

To account to stochasticity between trials for these predictions (which notably had a much larger impact likely due to the decreased constraints on rotamer optimization in these cases), this set of ΔΔG values was reported as an average of 50 trials. Significant outliers for overall binding affinity change from SARS-CoV to SARS-CoV-2 were called based on similar criteria to the individual mutations, except the z-score normalization was performed relative to all other interactions.

### Protein-ligand Docking Using Smina

The previous viral-human interactome screen by Gordon *et al.*^20^ reported 76 candidate drugs targeting the 332 human proteins. In order to further prioritize this list and search for drugs that share a binding site with the viral interactor, we performed protein-ligand docking. Among this list, 71 interaction-drug pairs involving 49 unique drugs that were amenable to docking were identified. To prep for docking, 3D structures for all ligands were first generated using Open Babel^113^ and the command:

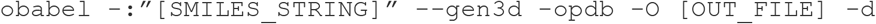

Protein-ligand docking was executed using smina^114^ with the following parameters. The autobox_ligand option was turned on and centered around the receptor PDB file with an autobox_add border size of 10 Å. The exhaustiveness was set to 40 to increase the number of independent stochastic sampling trajectories and increase the likelihood of identifying a global minimum. The num_modes was set to 1000 to so that a multitude of lower ranked models could be compared for internal purposes. To reduce real wall time each docking process was run using 5 CPU cores, although it should be noted runtime is affected in a strictly linear manner and CPU cores has no impact on net CPU time. Finally, each protein-ligand docking command was repeated 45 times with a unique seed on each interaction. The final smina command used was as follows:

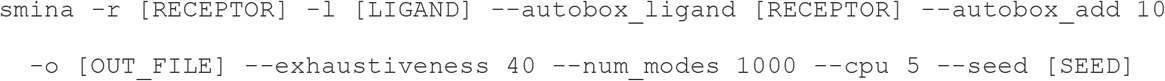

Multiple trials for the same docking problem were performed in order to saturate the ligand binding search space as thoroughly as possible. Practically speaking this batched docking approach was only taken so that time spent docking would cycle through each protein-drug pair equally, thus producing results from each pair at an even rate. The net effect is merely as if one docking process had been run for each protein-drug pair with exhaustiveness parameter set to ~2,000. It should be noted that this level of sampling was overkill relative to common practices and usually a single run with exhaustiveness ranging from 30-50 would be suffifient^114^. Docked conformation from all iterations on each protein-drug pair were compiled into a final set of up to 10 of the best scoring poses. To retain poses that cover different low-energy binding sites, these poses were selected such that the center of mass of each docked pose added must be at least 1 Å away from the center of mass of any of the higher ranked poses. All results described in this manuscript are reported based on only the top ranked pose for each protein-drug pair. Protein residues involved in the drug binding site were annotated using the same criteria used to define interface residues. Of note, to include ligand molecules as part of the NACCESS solvent accessible surface area calculations, the Record Type for all ligand atoms must be manually changed from HETATM to ATOM.

## Supplemental Table Headings

**Supplemental Table 1. List of ECLAIR predicted interface residues**.

**Supplemental Table 2. List of guided docking annotated interface residues**.

**Supplemental Table 3. List of human population variants reported by gnomAD**.

**Supplemental Table 4. List of sequence divergences between SARS-CoV and SARS-CoV-2**.

**Supplemental Table 5. Enrichment for sequence variation on SARS-CoV-2-human interfaces**.

**Supplemental Table 6. Enrichment for individual disease terms in human interactors of SARS-CoV-2**.

**Supplemental Table 7. Predicted ΔΔG between SARS-CoV and SARS-CoV-2 versions of all docked interactions**.

**Supplemental Table 8. Predicted ΔΔG impact of all human population variants at the interface**.

**Supplemental Table 9. List of all predicted drug-target binding sites**.

---

Figure 4. Drug Docking and Prioritization of SARS-CoV-2-Human Interaction Inhibitors

**Supplemental Figure 1.**
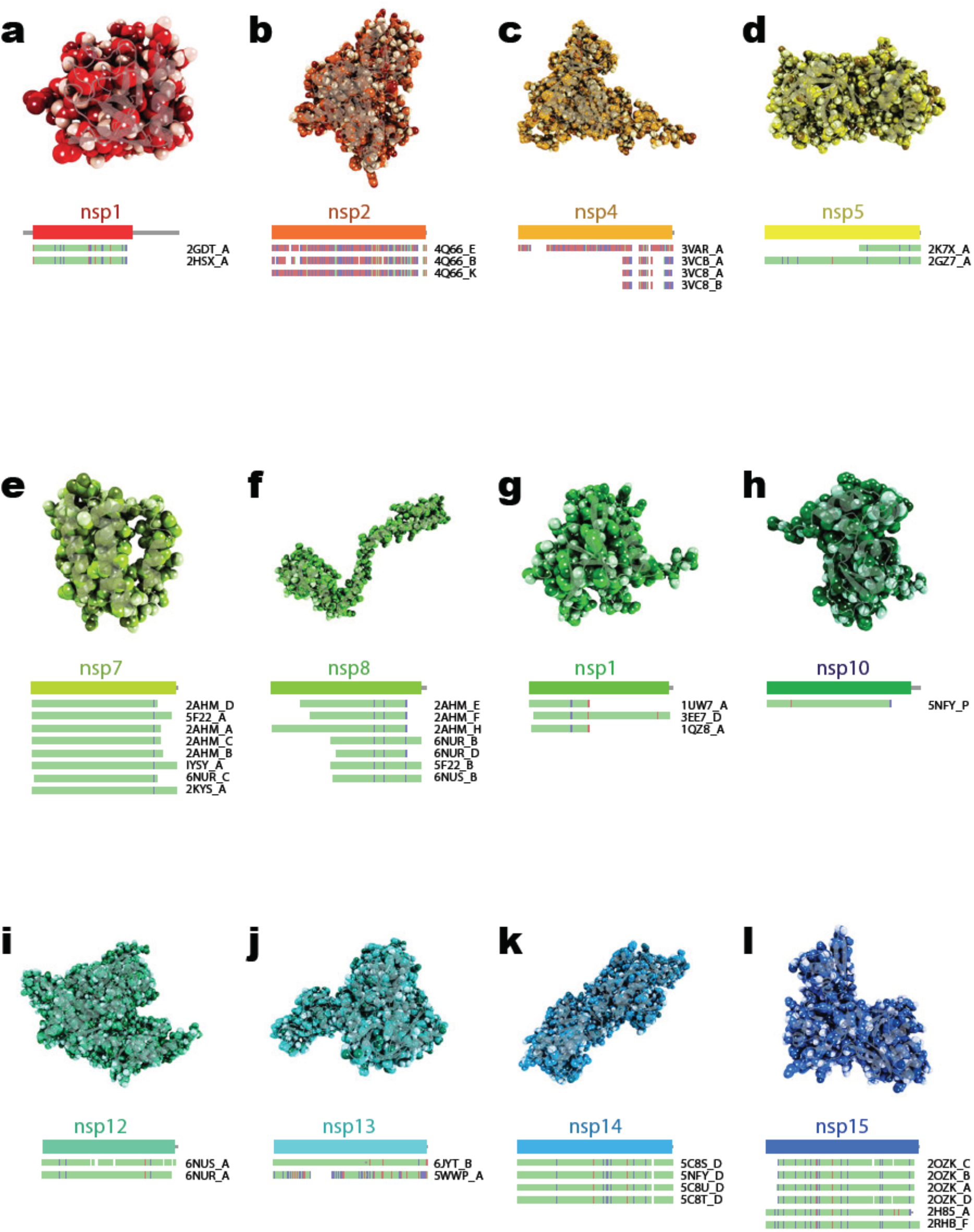
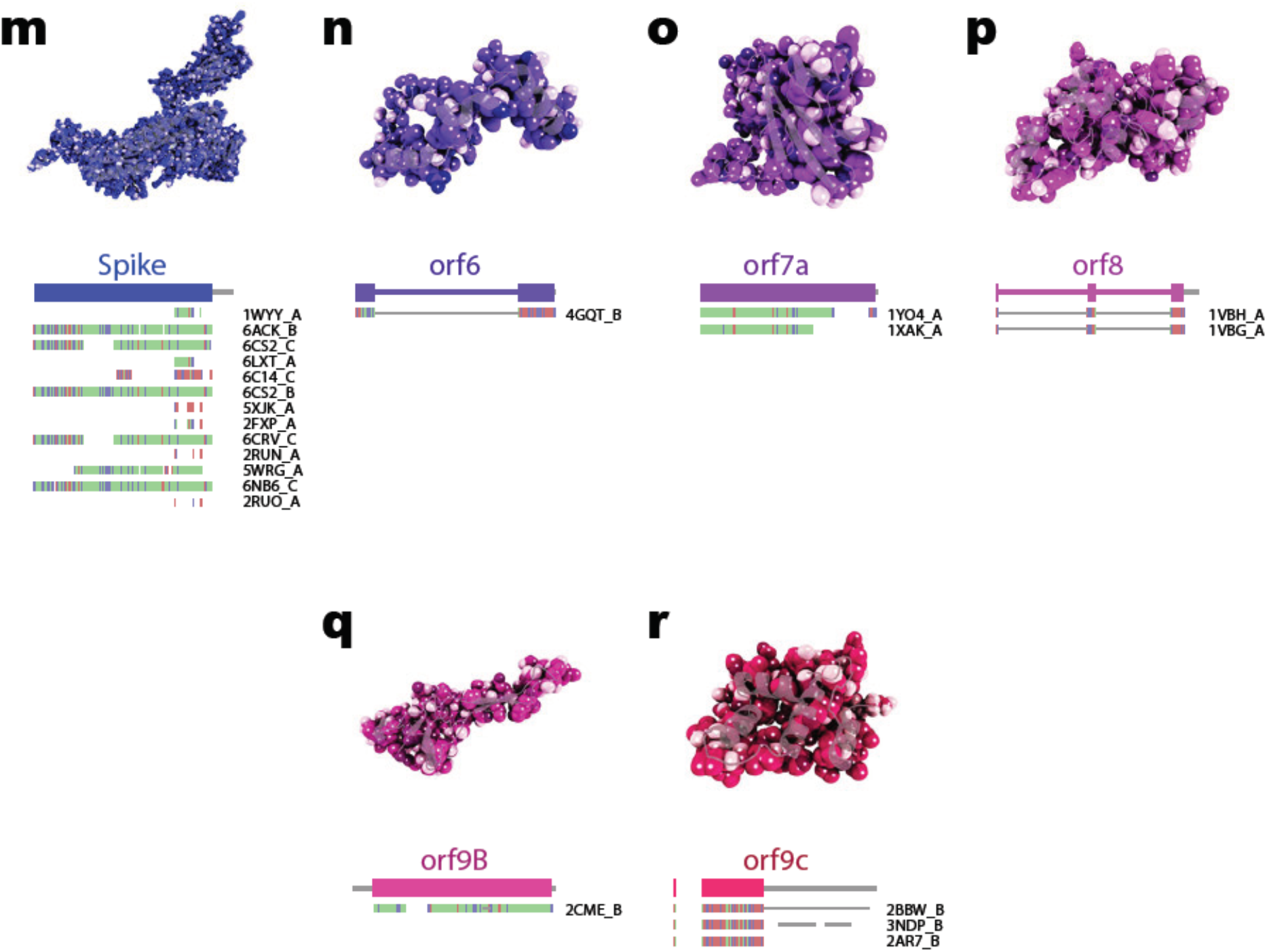
Homology modeling for SARS-CoV-2 proteins. **a-r,** Homology models for 18 SARS-CoV-2 proteins amenable to homology modeling. The first bar under each model indicates the overall coverage of the SARS-CoV-2 protein. The underlying bars indicate template utilization across the structure. The template bars are colored based on the alignment between the template and the corresponding SARS-CoV-2 where green indicates identical sequence, blue indicates a mismatch with positive BLOSUM substitution score, and red indicates a mismatch with a negative BLOSUM substitution score.

**Supplemental Figure 2.**
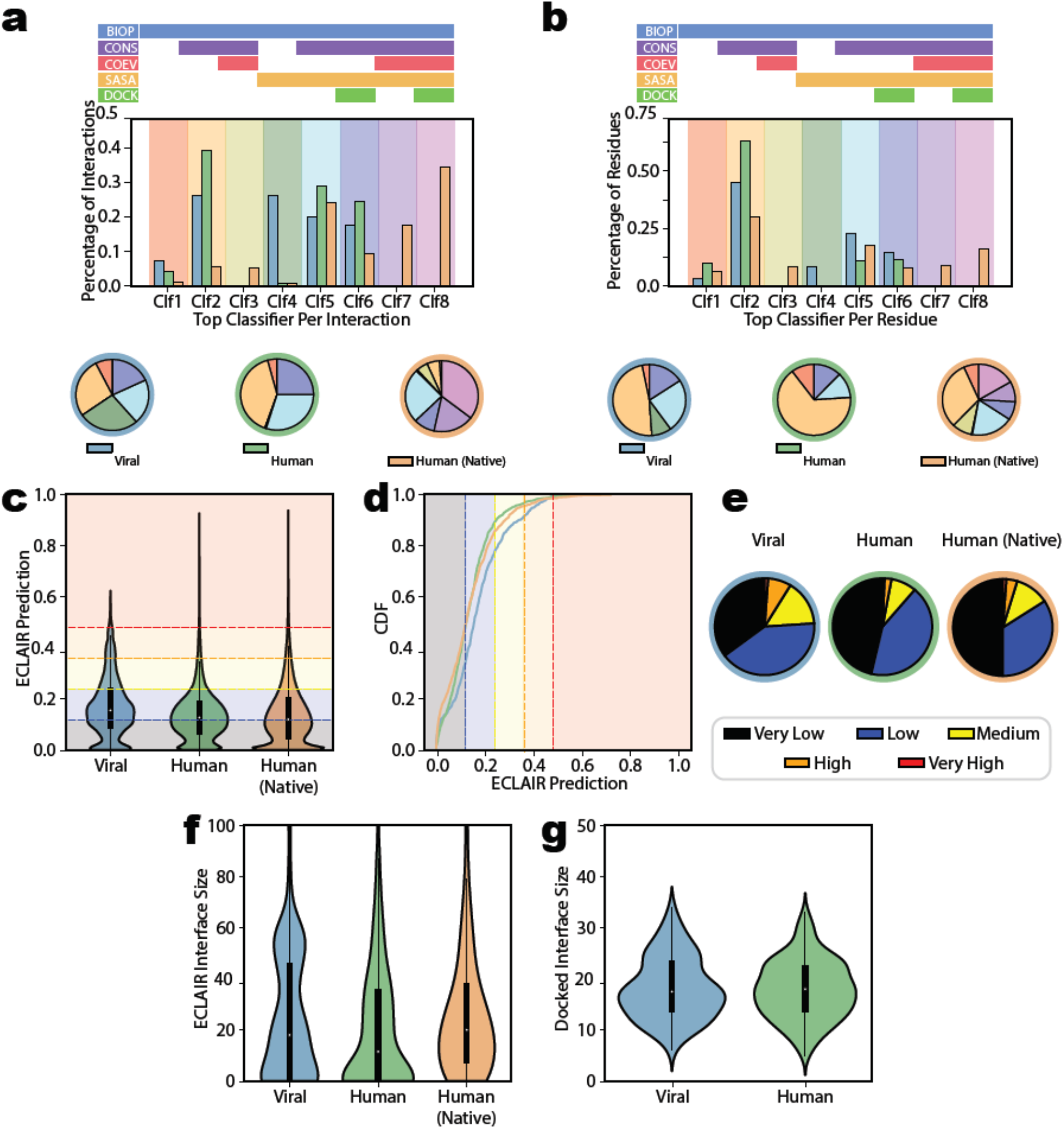
Summary of ECLAIR and guided docking interface predictions. **a,** Breakdown of best ECLAIR classifier used in interface prediction per interaction. The top section indicates which features are used by each classifier. The total breakdown of classifier usage for prediction of viral interfaces (blue) and human interfaces on human-viral (green) or native human-human (orange) interactions are shown either in bar plot (middle) or as pie charts (bottom). **b,** Identical breakdown, but reported at a per-residue rather than per-interaction level. **c, d,** Summary of ECLAIR prediction scores across viral, human, and native human-human interfaces presented as either a raw distribution or a cumulative density respectively. Plots are colored based on corresponding ECLAIR confidence bins in each range of the plot. **e,** Breakdown of the overall ECLAIR confidence bin representation among the three classes. Overall, ECLAIR classification predictions were generally similar between the three classes of interface, although a higher fraction of predictions made on viral interfaces were made with high or medium confidence. **f,** Distribution of the size of all interfaces defined by ECLAIR compared among the three classes. **g,** Distribution of viral or human interfaces defined by guided docking.

**Supplemental Figure 3.**
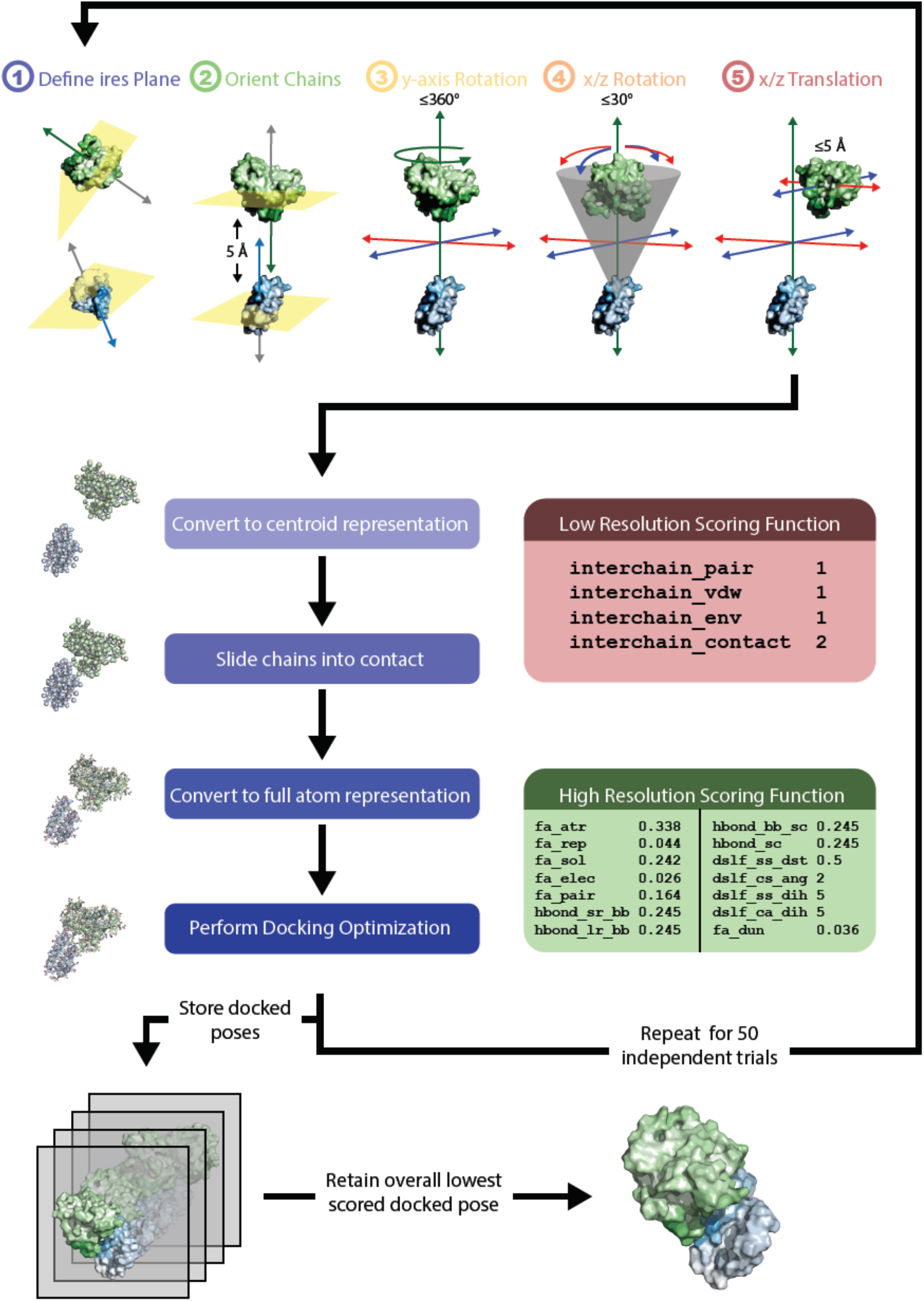
Visual representation of the guided docking protocol used. For each SARS-CoV-2-human interaction with 3D structure available for both proteins, 50 independent guiding docking trials were used to select a final docked configuration. The structure for the viral protein is colored from white to blue with darker blue corresponding to higher ECLAIR prediction. The structure for the human protein is colored similarly using a green to white gradient. Initial semi-random docked configurations were generated using five steps. First a plane separating ECLAIR predicted likely interface from likely non-interface residues was drawn to divide each protein. Second, the two protein chains were separated 5 Å apart on the y-axis using the previously defined plane to orient the likely interface sides of each protein towards each other. Third, the human protein was randomly rotated up to 360° along the y-axis to sample different orientations of the two interfaces relative to each other. Fourth, the human protein was randomly rotated up to 30° along the x- and z-axes with the point of rotation centered on the viral protein. Fifth a random translation up to 5 Å was applied to the human protein along the x- and z-axes. The first two steps constitute the ECLAIR-based guiding of the docking space. The last three steps serve to randomly perturb the initial docking configurations to sample the space near the ECLAIR predicted interface. After this initial perturbation docking is performed using a combination of low resolution and high-resolution scoring. During the low-resolution scoring, the proteins are initially converted to a centroid representation and slid into contact. During the high-resolution scoring, the proteins are converted back to a full-atom representation and the contact and side-chain packing is optimized. The best (lowest) scored docked pose is retained and used for docked interface definition.

**Supplemental Figure 4.**
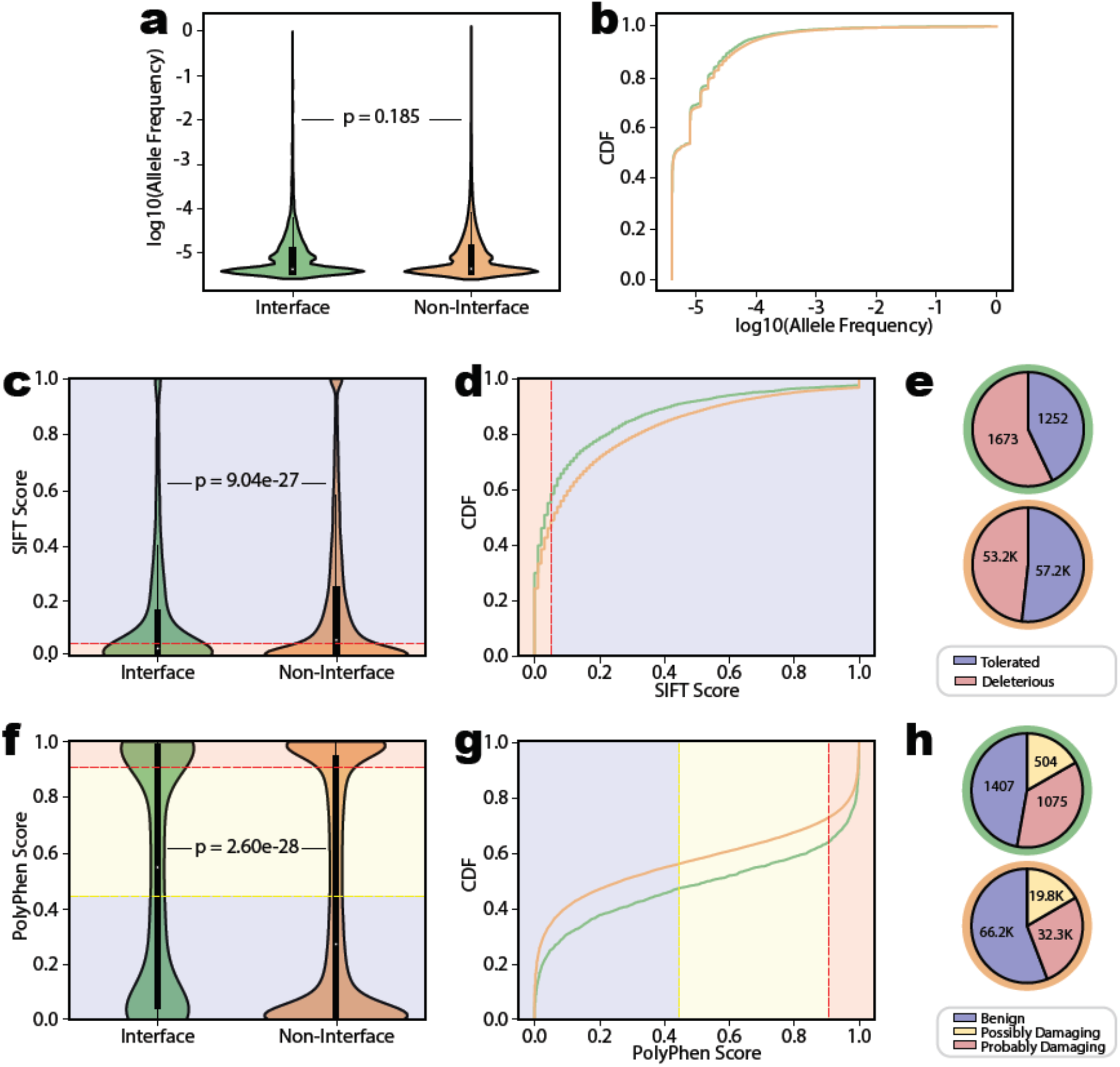
Summary of human population variant frequency and deleteriousness. **a, b,** Summary of allele frequency for human population variants either on or off the predicted human-viral interface presented as either a raw distribution or a cumulative density respectively. Variants in either category had roughly identical allele frequency distributions. **c, d,** Summary of the SIFT deleteriousness score for human population variants either on or off the predicted human-viral interface presented as either a raw distribution or a cumulative density respectively. Plots are colored based on the split between SIFT tolerated and deleterious categories. **e,** Pie chart breakdown of these categories. Pie char outlines distinguish interface (green) from non-interface (orange). Population variants on the interface were significantly more likely to be classified deleterious. **f, g,** Summary of the PolyPhen deleteriousness score for human population variants either on or off the predicted human-viral interface presented as either a raw distribution or a cumulative density respectively. Plots are colored based on the split between PolyPhen benign, possibly damaging, and probably damaging categories. **e,** Pie chart breakdown of these categories. Pie char outlines distinguish interface (green) from non-interface (orange). Population variants on the interface were significantly more likely to be classified probably damaging.

